# Smells like γ-synchrony: Insula-prefrontal communication depends on γ-synchrony and supports modality-specific changes in behavioral strategies

**DOI:** 10.64898/2026.08.06.743126

**Authors:** Lara L. Hagopian, Aarron J. Phensy, Austin J. Gallagher, Vikaas S. Sohal

## Abstract

The prefrontal cortex is critical for many aspects of flexible behavior, but how it interacts with other brain regions to perform this function remains largely unknown. The insula is bidirectionally coupled with prefrontal cortex and known to be necessary for many aspects of cognition. Here we examined how the medial insular cortex (mIC) and medial prefrontal cortex (mPFC) interact to promote flexible behavior. Disrupting mIC-mPFC connectivity interferes with the ability of mice to learn shifts from texture cue-based behavioral strategies to odor-based ones (but not vice-versa). Using genetically encoded voltage indicators, we find a corresponding increase of in-phase gamma-frequency synchronization between mIC and mPFC parvalbumin-expressing inhibitory neurons during texture-to-odor shifts. Finally, we confirmed that optogenetically perturbing this synchronization disrupts texture-to-odor (but not odor-to-texture) shifts. These results establish a critical role for gamma synchronization in insula-prefrontal communication, and show that this communication plays a sensory modality-specific role in flexible behavior.

## INTRODUCTION

A key factor underlying survival in a dynamic environment is the ability to develop behavioral strategies that utilize information from multiple sensory modalities and flexibly adapt when the environment changes. The prefrontal cortex is believed to play a key role in this type of cognitive flexibility ^1^. Deficits in cognitive flexibility, measured by set shifting tasks such as the Wisconsin Card Sorting Task (WCST), are major contributors to disability in schizophrenia and related neuropsychiatric conditions ^2,3^, underscoring the importance of identifying mechanisms through which the prefrontal cortex can flexibly change how it uses sensory information to guide behavior.

Presumably, the ability of the prefrontal cortex to flexibly utilize competing streams of incoming sensory information involves interactions with other brain regions. The insula is a likely candidate for this based on several factors. First, the insula has extensive bidirectional connectivity with many prefrontal regions involved in cognitive flexibility, including the medial prefrontal cortex (mPFC) in rodents ^4,5^. Second, lesion studies in humans implicate the insula in multimodal sensory integration generally ^6^ and in set-shifting specifically ^7^. Consistent with this, dysfunction of the insula has been suggested to contribute to the development of schizophrenia ^8–11^. Third, in addition to its roles in interoception, the insula is known to process several types of sensory information, particularly taste and olfactory information ^12,13^, and to facilitate learning about new tastes ^14^. Insular neurons also respond to sensory cues that predict food rewards and may serve to encode relationships between those cues, associated actions (e.g., reward consumption), and the interoceptive consequences of those actions ^15^.

This raises the question – how do the prefrontal cortex and insula interact to support cognitive flexibility? The insula may be involved in cognitive flexibility generally, or it may provide prefrontal circuits with information related to certain sensory modalities. Further, if the prefrontal cortex and insula do interact, what circuit processes might mediate this interaction? Several studies from our laboratory have shown that gamma-frequency (∼40 Hz) synchronization among parvalbumin-expressing inhibitory neurons (PV INs) in the mPFC plays a critical role in cognitive flexibility, specifically during a rule-shifting task, in which mice learn to find hidden food rewards using odor or texture cues ^16–20^. In this context it is unclear whether gamma synchrony might extend beyond prefrontal cortex to the insula, and if so, whether it could facilitate any interactions between these regions that are necessary for cognitive flexibility? Investigating the possibility that gamma activity in the prefrontal cortex and insula synchronizes is important; prior work from our lab and others has established that gamma synchronization is essential for cognitive flexibility, but has not fully resolved its function. In particular, a long-standing hypothesis, ‘Communication Through Coherence’ (CTC) posits that interregional gamma synchrony facilitates coordinated interactions between those regions ^21,22^.

Here using optogenetic inhibition, we confirm that projections from the medial insular cortex (mIC) to the mPFC are essential for cognitive flexibility, specifically when animals learn to shift from rules based on texture cues to ones based on odor cues. Notably, mIC-mPFC connections are not required for the opposite type of shift or for de novo learning of odor-based rules. Then, using genetically encoded voltage indicators (GEVIs) we confirm that new patterns of gamma synchrony between PV INs in the mPFC and mIC emerge during texture to odor shifts; furthermore, optogenetically disrupting these patterns is sufficient to disrupt the texture to odor shifts. Together these results reveal that long-range gamma synchronization contributes to insula-prefrontal communication which plays a modality-specific role in cognitive flexibility.

## RESULTS

To examine the role of mIC to mPFC communication in cognitive flexibility, we optogenetically inhibited the cell bodies of mIC-to-mPFC (mIC-mPFC) projection neurons during a rule shifting task. Wild-type (WT) mice were injected with AAVrg-EF1a-Cre in the mPFC and either AAV5-Ef1a-DIO-eNpHR-eYFP (‘eNpHR+’, experimental group) or AAV5-Ef1a-DIO-eYFP (‘eYFP+’, control group) in the mIC. Optical fibers were implanted bilaterally in the mIC (Fig 1a-b).

**Figure 1:**
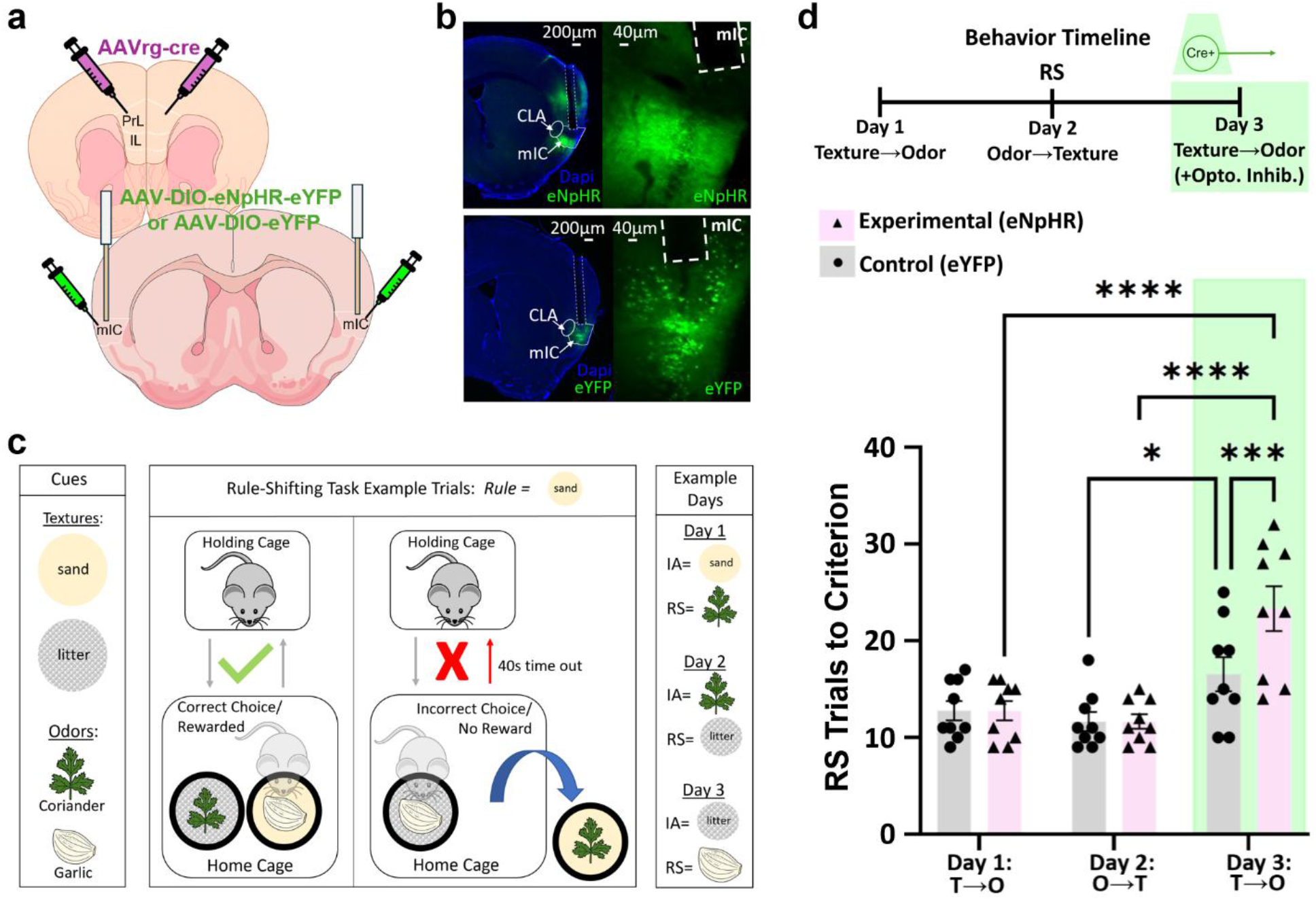
Inhibition of mIC-to-mPFC projection neurons impairs texture-to-odor rule shifts. **a,** Schematic: AAVrg-cre was bilaterally injected into mPFC, The mIC received bilateral injections of either AAV-DIO-eNpHR-eYFP or AAV-DIO-eYFP plus fiber optic implants (CLA = claustrum). **b,** Example histological images showing virus expression and fiber localization (Top: eNpHR-eYFP; Bottom: YFP). **c,** Example of the rule shift task. Left: the four cues. Middle: an example of a correct and incorrect trial (the correct bowl is indicated by the location of the sand cue). Right: an example of how the rules / correct cue evolve across 3 consecutive days of rule shifting. The rule from the RS phase on one day is the IA rule on the following day. **d,** Top: Experimental timeline. Cell bodies of mIC-mPFC neurons receive optogenetic inhibition during each RS trial on Day 3. Bottom: there is a significant deficit in the learning of the texture to odor (‘T→O’) shift on Day 3 in experimental (eNpHR+) mice receiving optogenetic inhibition (indicated by the green shading). * p < 0.05. *** p < 0.001. **** p < 0.0001.

### Inhibiting mIC-mPFC projection neurons disrupts texture-to-odor rule shifts

After 8-9 weeks, mice were run on the rule shifting task for 3 days. In this task, mice are presented with two bowls on each trial. Each bowl is filled with a different textured medium (either sand or litter) and scented with a different odor (either garlic or coriander). The pairing of odor and texture cues within the same bowl varies randomly from trial to trial. Mice had to first learn a rule based on one modality (e.g., ‘dig in the sand-containing bowl’), then shift to a rule based on the other modality (e.g., ‘dig in the garlic-containing bowl’). These two rules represent the initial association (IA) and rule shift (RS) phases of the task, respectively (Figure 1c; Methods). Mice were run for three days on this task comprising both IA and RS phases (Day 1: texture to odor shift; Day 2: odor to texture shift; Day 3: texture to odor shift). Mice received optogenetic inhibition on Day 3, specifically during RS trials to test the role of mIC-mPFC input on shifts to odor cues (Fig. 1c-d). Optogenetic inhibition was turned off once mice were placed back in the holding cage during intertrial intervals (including extended timeouts after error trials).

We observed a significant deficit in RS performance, as measured by the number of trials needed to reach the learning criterion (8/10 consecutive trials correct), during Day 3 for experimental (eNpHR+) mice compared to Day 1 (p<0.0001) and Day 2 (p<0.0001). Experimental (eNpHR+) mice also have significantly worse performance than control (eYFP+) mice on Day 3 (p=0.0001). Control (eYFP+) mice showed no significant difference in RS performance between Day 1 and Day 3 (p=0.11). Of note, control mice did have a slight, but significant, deficit in performance on Day 3 compared to Day 2 (p=0.03), although again the performance of experimental (eNpHR+) mice on Day 3 was markedly worse. The slightly worse performance of control mice on Day 3 relative to Day 2 (but not Day 1), may reflect a difference in the difficulty of texture to odor shifts, compared to odor to texture shifts, although as will be shown below, this difference did not consistently occur in subsequent experiments.

### mIC-mPFC projection neurons are not required for odor-to-texture shifts or learning initial associations based on odor cues

The preceding results show that mIC-mPFC projection neurons are necessary for shifts from texture to odor-based rules. This raises the question of whether these neurons are necessary for rule shifts in general (e.g., for odor-to-texture shifts) or for detecting or processing odor cues more generally? To answer these questions, we conducted two new sets of experiments, analogous to the preceding ones, in which we optogenetically inhibited mIC-mPFC projection neurons either when mice were shifting from odor to texture-based rules, or when they learned an initial odor-based rule.

Wild-type mice were injected with viruses and implanted with optical fibers as described above (Fig. 1a-b). To examine the effects of inhibiting mIC-mPFC neurons on odor-to-texture shifts, mice were run on the rule shift task over three days: Day 1, odor to texture shift; Day 2, texture to odor shift; Day 3, odor to texture shift. Mice in this experiment received optogenetic inhibition on Day 3 during RS trials (Fig. 2a). As before, optogenetic inhibition was turned off when mice were in the holding cage during intertrial intervals. We observed no significant difference in performance (the number of RS trials needed to reach the learning criterion) of experimental (eNpHR+) mice on Day 3 compared Days 1 and 2. There also was no significant difference in performance on Day 3 between experimental (eNpHR+) and control (eYFP+) mice. This suggests that, in contrast to shifts to odor-based rules, mIC-mPFC projection neurons are not necessary for shifts to texture-based rules.

**Figure 2:**
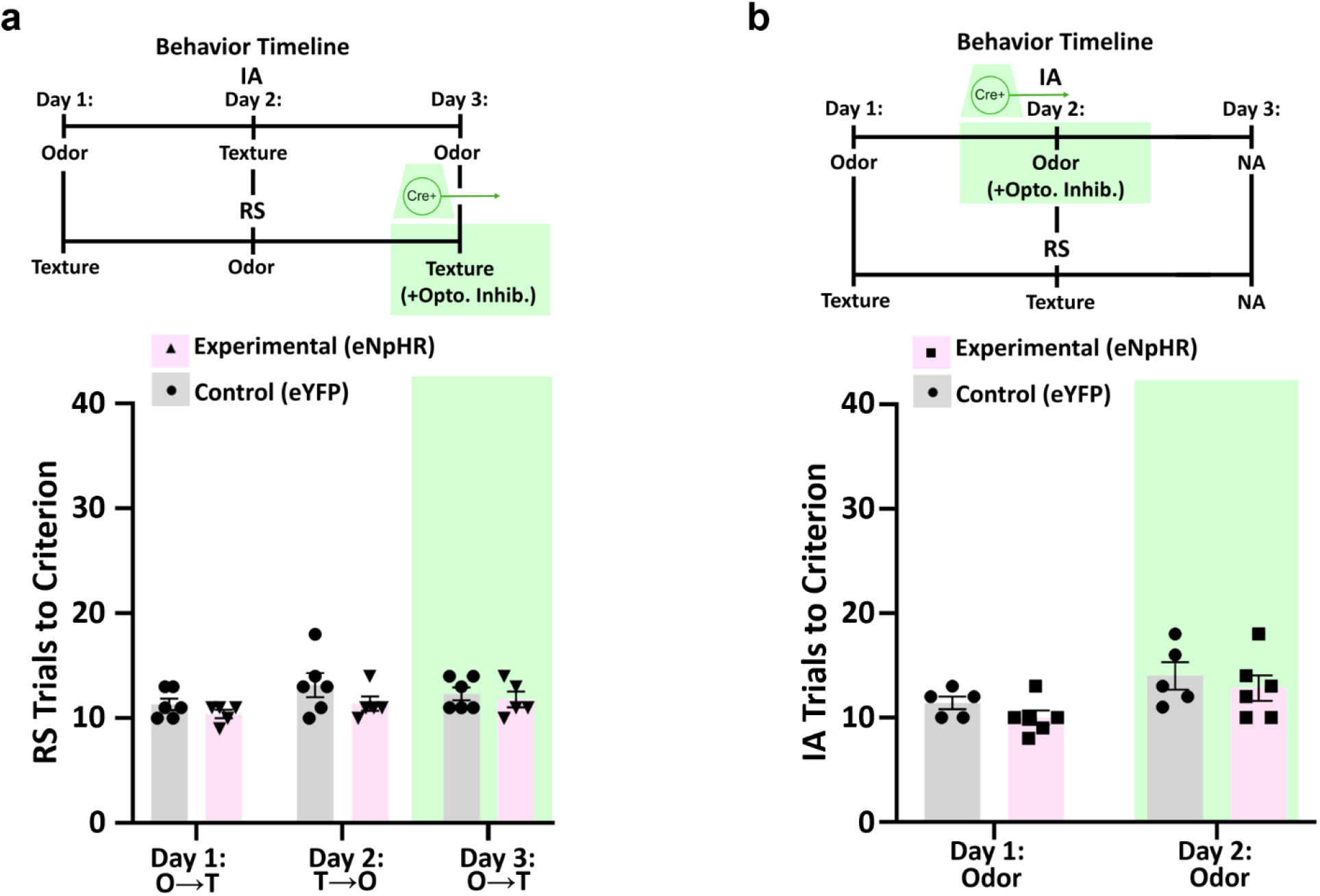
Inhibiting Insula-to-mPFC projection neurons does affect shifts to texture-based rules or the initial learning of odor-based rules. **a,** Top: Behavioral timeline. mIC-mPFC neurons receive optogenetic inhibition during each RS trial on Day 3. Bottom: RS performance (number of trials needed to reach the learning criterion) across days for odor to texture (O→T) and texture to odor (T→O) shifts in experimental (eNpHR+, pink bars) and control (eYFP+, gray bars) mice. Green shading indicates the delivery of optogenetic inhibition on Day 3. **b,** Top: Behavioral timeline. mIC-mPFC neurons receive optogenetic inhibition during each trial when learning an initial odor-based association on Day 2. Bottom: there is no significant difference in learning of the odor rule (number of trials needed to reach the learning criterion) for experimental (eNpHR+) vs. control (eYFP+) mice on Day 2, or for experimental mice on Day 1 vs. Day 2.

To evaluate whether inhibiting mIC-mPFC neurons more generally disrupts the ability of mice to detect, learn from, or make decisions based on odor cues, mice were run for two days on a version of the task involving just a single association between an odor cue and food reward: Day 1, IA based on one odor cue; Day 2, IA based on the other odor cue. On Day 2, mice received optogenetic inhibition during IA trials (Fig. 2b). Again, optogenetic inhibition was turned off when mice were in the holding cage during intertrial intervals. There was no significant change in performance (the number of trials needed to reach the learning criterion) for experimental (eNpHR+) mice on Day 2 relative to Day 1, or between experimental and control (eYFP+) mice on Day 2.

### Inhibiting mIC-mPFC axon terminals is sufficient to disrupt shifts to odor-based rules

Our previous results (Fig. 1) show that mIC-mPFC projection neurons are necessary for texture to odor shifts. However, given the extensive connectivity between the insula and other brain regions ^4^, it is unclear whether this specifically reflects the projections of these neurons to the mPFC, their projections to other brain regions, or local effects within the mIC. To directly test the role of mIC projections to the mPFC, we delivered optogenetic inhibition to mIC-mPFC axon terminals when mice learned shifts to odor-based rules.

Wild-type (WT) mice were again injected with AAVrg-EF1a-Cre (‘retro-cre’) in the mPFC and either AAV5-Ef1a-DIO-eNPHR-eYFP (experimental group, ‘eNpHR+’) or AAV5-Ef1a-DIO-eYFP (control group, ‘eYFP+’) in mIC (bilaterally). Optical fibers were implanted bilaterally in mPFC (Fig 3a-b). Mice were run for three days on the rule shifting task: Day 1, texture to odor shift; Day 2, odor to texture shift; Day 3, texture to odor shift. Mice received optogenetic inhibition during RS trials on Day 3 (Fig. 3c-d). As before, optogenetic inhibition was turned off when mice were in the holding cage during intertrial intervals. Optogenetic inhibition disrupted the learning of shifts to odor-based rules, as evidenced by a significant increase in the number of RS trials needed to reach the learning criterion for experimental (eNpHR+) mice on Day 3 compared to Days 1 (p=0.0019) and 2 (p=0.0012). Experimental mice also performed significantly worse than control (eYFP+) mice on Day 3 (p=0.0086). There was no significant difference in RS learning for control (eYFP+) mice on Day 3 vs. Days 1 (p=0.58) or 2 (p=0.67).

**Figure 3:**
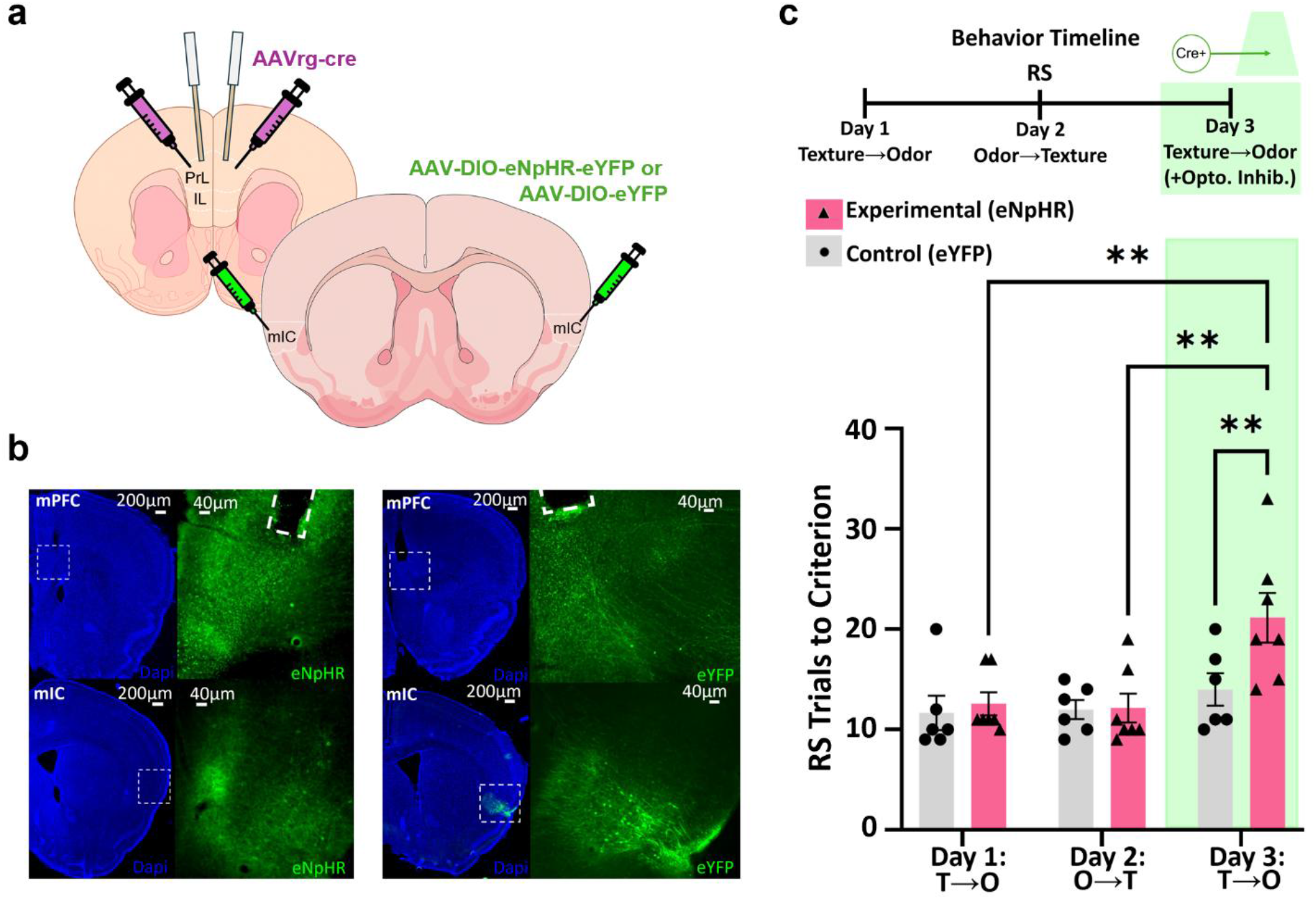
Inhibiting Insula-to-mPFC projections disrupts shifts to odor-based rules. **a,** Schematic of bilateral virus injections (into mPFC and mIC) and fiber implants (in mPFC). **b,** Example images showing viral expression and implant locations (left: experimental mouse expressing eNpHR-eYFP, right: control mouse expressing eYFP only). **c,** Top: Behavioral timeline. mIC-to-mPFC terminals receive optogenetic inhibition during RS trials on Day 3. Bottom: Optogenetic inhibition causes a significant deficit in RS performance (number of trials needed to reach learning criterion) in experimental mice on Day 3. ** p < 0.01.

### mIC and mPFC PV interneurons differentially synchronize during texture-to-odor vs. odor-to-texture shifts

The preceding results establish that mIC-mPFC communication is essential for learning rule shifts from texture to odor. As described earlier, previous work from our laboratory has shown that within the mPFC, gamma-frequency synchronization involving parvalbumin-expressing inhibitory neurons (PV INs) plays a key role in rule shift learning. This raises the question: does such gamma-synchrony extend to the mIC, and if so, is it associated with particular aspects of rule shift learning? To address this question, we used Transmembrane Electrical Measurements Performed Optically (TEMPO) to quantify gamma-frequency (30-50Hz) synchronization between mIC and mPFC PV INs. TEMPO uses genetically encoded voltage indicators (GEVIs) to measure membrane voltage dynamics on fast time scales, comparable to local field potential, but with cell type-specificity ^23,24^. We injected PV-Flp mice in both the mPFC and mIC with AAV1-CAG-fDIO-Ace2N-4AA-mNeon to express the GEVI, Ace-mNeon, in PV INs, as well as AAV-syn-tdTomato to generate a control fluorophore signal. We implanted optical fibers for photometry unilaterally in both the mIC and mPFC (Figure 4a-b).

**Figure 4:**
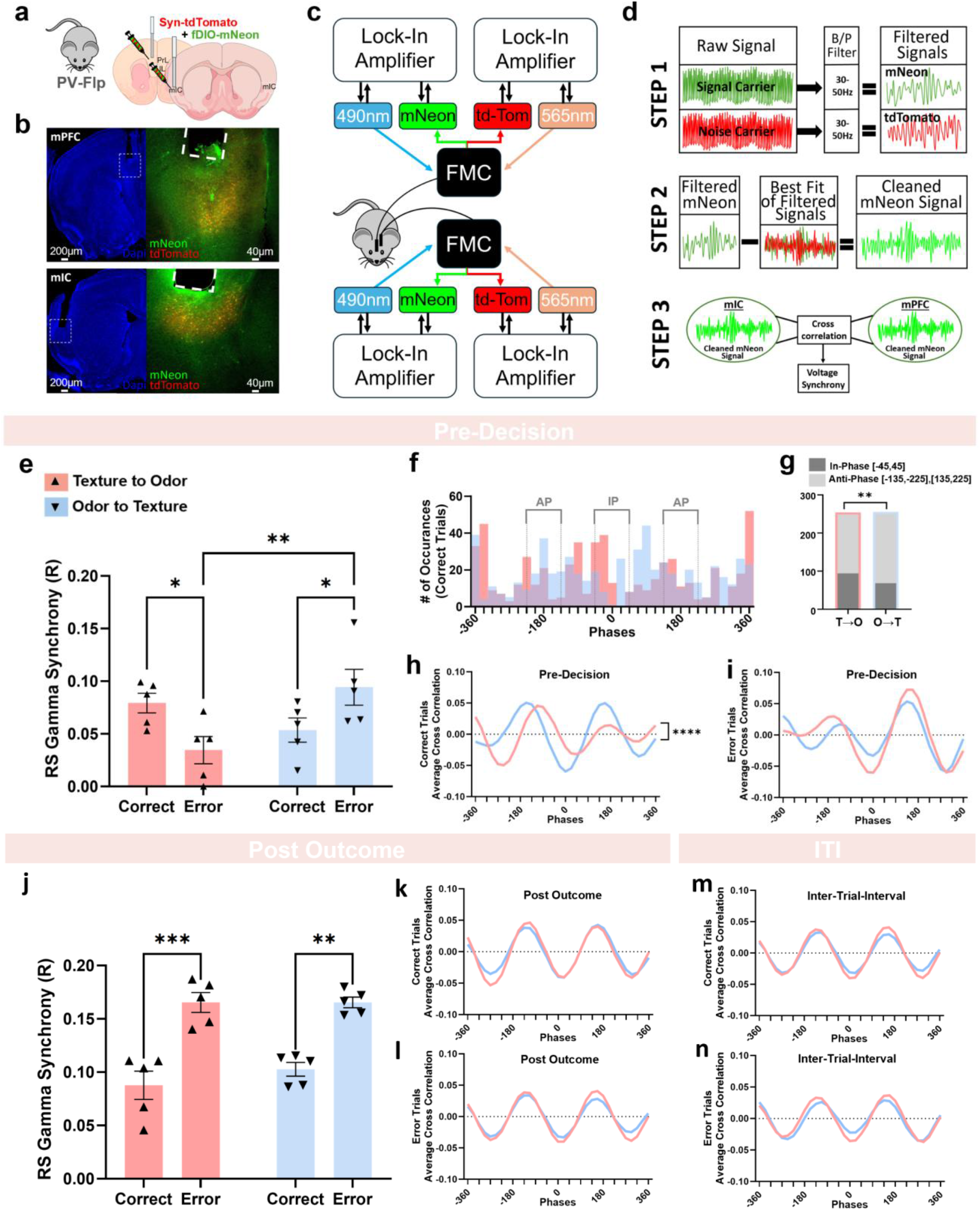
Gamma synchrony between mIC and mPFC PV interneurons changes depending on whether mice use odor or texture-based strategies. **a,** We injected a 2:1 mixture of AAV1-CAG-fDIO-Ace2N-4AA-mNeon (‘fDIO-mNeon’) and AAV2-Syn-tdTomato (‘Syn-tdTomato’) and implanted optical fibers unilaterally into mIC and mPFC. **b,** Example image showing viral expression and implant locations (Top: mPFC; Bottom: mIC). **c,** Schematic of hardware for TEMPO recordings. Dual-site fiber photometry was used to measure mNeon and tdTomato signals. ‘FMC’ = filter mini-cube. Lock-in amplifiers modulated and demodulated excitation and emission light, respectively. **d,** Gamma band activity is extracted by filtering raw fluorescent signals, first between 1-100Hz, then from 30-50Hz. After each filtering step, the best fit between mNeon and tdTomato signals is used to remove shared noise components. Finally, cross correlation is used to quantify synchrony between PV IN activity in mPFC and mIC. **e-i:** mIC-mPFC PV IN gamma synchrony (peak cross-correlation) during pre-decision periods (1-3 sec preceding each dig). **e,** Pre-decision gamma synchrony on correct vs. error trials during texture to odor or odor to texture shifts. Synchrony is significantly different on correct and error trials for texture to odor vs. odor to texture shifts, and on error trials for shifts to odor vs. texture. **f,** Distributions of mIC-mPFC phase differences (i.e., locations of the peak cross correlation) from −360 to +360 degrees (AP: Anti-Phase; IP: in-phase). **g,** There is a significant difference in the fraction of timepoints which are in-phase vs. anti-phase for texture to odor vs. odor to texture shifts. **h-i,** averaged cross correllograms are significantly different for texture to odor vs. odor to texture shifts on correct (h) but not error (i) trials. **j-l**, mIC-mPFC PV IN gamma synchrony (peak cross-correlation) during post outcome periods (1-3sec preceding each dig). **j**, Similar to (e) but for post outcome periods (trial outcome to trial end). In-phase mIC-mPFC PV IN gamma synchrony is significantly difference between correct and error trials for both texture to odor and odor to texture shifts. There is no significant difference between odor and texture shifts for either correct or error trials. **k-l**, similar to h-i but for post-outcome periods. **m-n,** Similar to h-I but for intertrial intervals (ITIs). * p < 0.05. ** p < 0.01. ***p<0.001. **** p < 0.0001.

After waiting 4-5 weeks for viral expression, we ran mice on two days of rule shifting. Mice learned an odor-to-texture shift one day, and a texture-to-odor shift on the other day. We randomly counterbalanced whether each mouse started with odor or texture-based rules on Day 1. TEMPO was performed following our previously described methods ^17,18,20^. Briefly, we used 490nm and 565nm LEDs to excite Ace-mNeon and tdTomato, respectively, along with lock-in amplifiers to modulate LED output and demodulate measured fluorescence in order to minimize cross-talk between fluorophores and recording sites (Figure 4c; Methods).

We analyzed TEMPO signals using the approach described in ^20^. Briefly, we first applied a bandpass filter (1-100Hz) to raw fluorescence signals. Then, we used robust linear regression to estimate shared noise components between Ace-mNeon and tdTomato signals, and subtracted this shared component from the broadband Ace-mNeon signals. We filtered the resulting signals in the gamma (30-50Hz) band, then once again used robust linear regression between the gamma-band filtered Ace-mNeon and tdTomato signals to remove shared noise components and obtain a ‘cleaned’ Ace-mNeon signal for both mIC and mPFC PV INs. Finally, we computed the cross correlation between mIC and mPFC signals, and used the peak value of the cross-correlation between −45 and +45 degrees to specifically quantify in-phase gamma synchrony. We used this metric to compare synchrony during pre-decision (1-3s before the mouse starts to dig) and post-outcome (time between the outcome and trial end) periods, depending on whether the trial was a correct decision or error. We quantified synchrony separately for odor-to-texture vs. text-to-odor shifts (Figure 4e-i).

During the pre-decision period, we observed different levels of mIC-mPFC PVI in-phase gamma synchrony during shifts to odor vs texture-based rules (Figure 4e-g). During shifts to odor-based rules, correct trials have significantly greater in-phase gamma synchrony than error trials (p=0.02), but the opposite is true during shifts to texture-based rules (correct vs. error trial synchrony: p=0.04) (Figure 4e). During rule shifts, error trials primarily occur when mice have not yet learned the proper rule to use for decision making. Thus, the double dissociation we observed suggests that pre-decision mIC-mPFC PV IN in-phase gamma synchrony rises when mice correctly learn texture→odor shifts, whereas the opposite occurs during odor→texture shifts. Indeed, focusing specifically on conflict trials (when the IA and RS cues are located in opposite bowls) during early portions of the RS shift, mIC-mPFC PV IN in-phase gamma synchrony is significantly higher for odor→texture compared to texture→odor shifts (Supplemental Figure 2b). Together, these findings support a model in which mIC-mPFC coordination involves in-phase gamma synchrony between PV INs in both regions, and differs depending on the cue modality being used for decision making.

To examine the nature of mIC-mPFC PV IN gamma-frequency synchronization in greater detail, we plotted the distribution of pre-decision phase differences between the cleaned, 30-50Hz filtered Ace-mNeon signals from both regions during the final 8 correct trials of the rule shift. The phase difference was the location of the overall peak in the cross-correlogram, computed for lags from +/− 1 cycle (i.e., +/− 25 msec). During texture→odor shifts, there was a peak in the phase lag distribution centered around 0 degrees. By contrast, there was a noticeable paucity of phase lags near 0 degrees during odor→texture shifts, and a relative enrichment of lags corresponding to anti-phase synchronization (∼180 or −180 degrees; Figure 4f-g). Thus, mIC-mPFC PV IN in-phase gamma synchrony occurs more often on correct trials during texture→odor shifts than during odor→texture shifts, and when it occurs, it is stronger (i.e., associated with higher correlation amplitudes).

To visually illustrate this difference, we also plotted the averaged cross-correlogram (Figure 4h-i). During the pre-decision period of correct trials for odor→texture shifts (Figure 4h), the value of the cross-correlogram at zero phase shifts is strongly negative, consistent with predominately anti-phase synchronization. However, during texture→odor shifts, this value becomes closer to zero, consistent with a relative enrichment of in-phase synchronization during these types of shifts. By contrast, on error trials (Figure 4i) we see the opposite pattern: more anti-phase synchronization for texture→odor shifts, and a shift towards in-phase synchronization for odor→texture shifts.

To evaluate whether this cue modality-dependence is specific for particular aspects of the rule shifting task, e.g., decision-making vs. processing outcomes, we also examined gamma synchrony during the post-outcome period (Figure 4j-l). There was no difference in mIC-mPFC PV IN in-phase gamma synchrony between shifts to odor and shifts to texture for either correct or error trials. Notably, there was a significant increase in this synchrony on error trials compared to correct trials for both shifts to odor (p=0.0004) and shifts to texture (p=0.001; Figure 4j). To visually illustrate the similarity in patterns of post-outcome gamma synchronization during texture to odor vs. odor to texture shifts, we also plotted averaged cross-correlograms (Figure 4k-l). During the post-outcome period of correct (Figure 4k) or error (Figure 4l) trials, the cross-correlograms for odor→texture vs. texture→odor shifts are largely overlapping. A similar pattern was observed during the inter-trial-interval (ITI; Figure 4m-n). These results suggest that the cue modality-dependence of mIC-mPFC PV IN gamma-synchrony is specific for decision making during pre-decision period, and does not generalize to the processing of outcomes during post-outcome or ITI periods.

### In-phase mIC-mPFC PV interneuron gamma synchrony is necessary for shifts to odor-based rules

The preceding analyses confirm that when mice are making decisions, patterns of mIC-mPFC PV IN gamma synchrony differ depending on whether mice shift to odor vs. texture-based rules. During shifts to odor vs. texture, there is a relative enhancement of in-phase vs. anti-phase synchronization, respectively. This raises the question: are these differences in synchronization just markers of shifts in decision-making strategies, or do they causally contribute to those shifts?

We addressed this question using a method we have previously leveraged to evaluate the causal significance of different patterns of inter-regional interneuron synchronization ^17,25^. Specifically we delivered rhythmic (40Hz) trains of light flashes to optogenetically excite interneurons in both structures, but did so either in-phase (same pattern delivered to both sites, ‘IP’) or out-of-phase (pattern shifted 144 degrees between sites, ‘OOP’). As in our experiments using optogenetic inhibition, stimulation was delivered specifically during trial periods, and turned off during inter-trial intervals.

We targeted PV-interneurons using PV-Cre mice (experimental group) and used Cre-negative mice for controls. We injected AAV5-DIO-ChR2-eYFP bilaterally in the mPFC and mIC (Figure 5a-b) and implanted a dual optical fiber in mPFC as well as optical fibers in mIC bilaterally. While some virus likely spread to neighboring regions (e.g., claustrum), the positioning of the implant directly above mIC should concentrate PV IN stimulation within this region.

**Figure 5:**
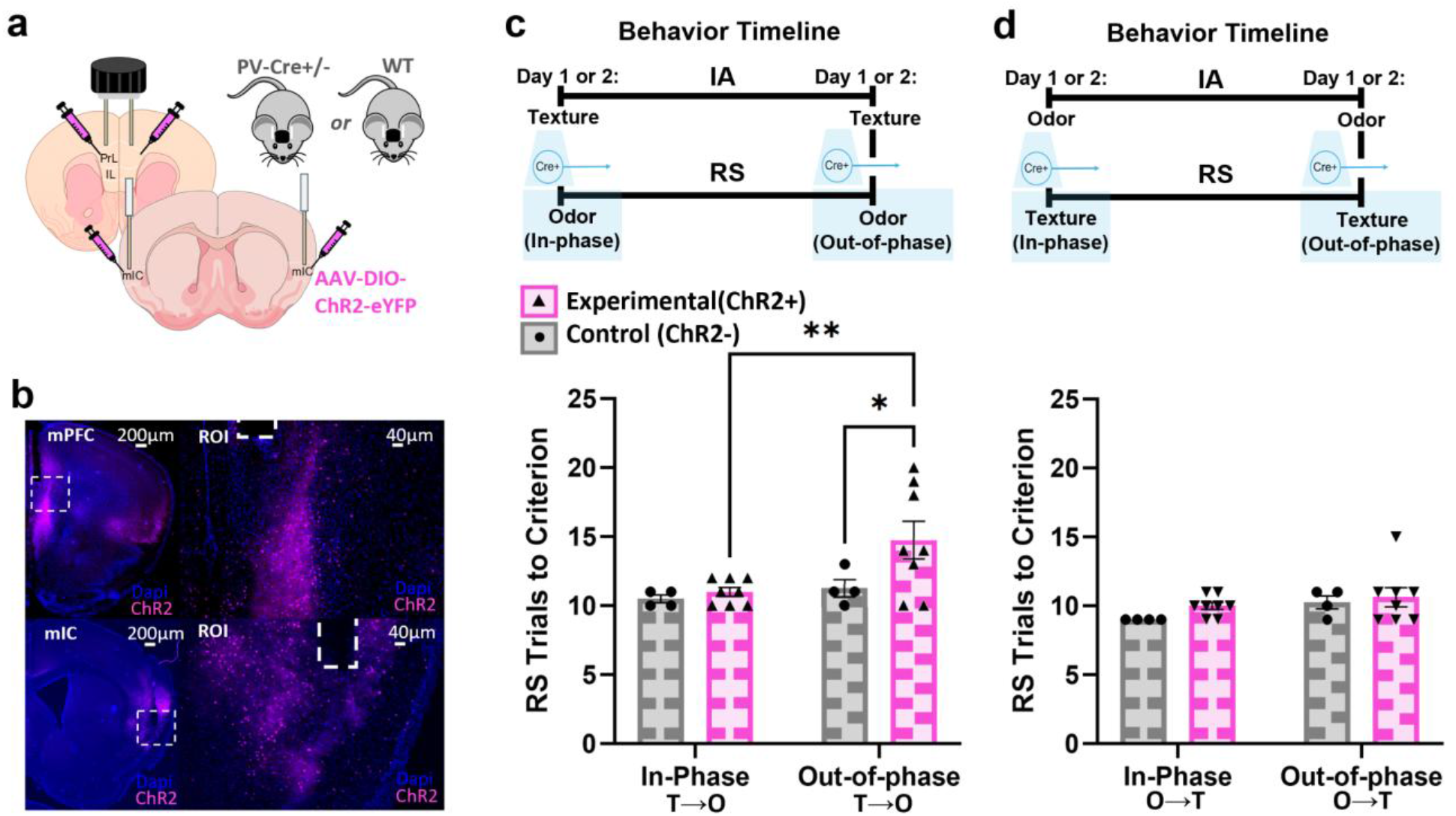
In-phase mIC-mPFC PV IN gamma synchrony is necessary for texture→odor shifts. **a,** Schematic of virus injection and optical fiber implants into bilateral mPFC and mIC. **b,** Example images showing virus expression and implant locations. **c,** Top: experimental timeline for optogenetic stimulation during texture→odor shifts. PV INs in mPFC and mIC are stimulated during each RS trial (in-phase one day, out-of-phase the other). Bottom: there is a significant increase in the trials needed to reach the learning criterion on the day with out-of-phase stimulation, compared to the in-phase day in experimental (PV-Cre) mice; there is also a significant increase in trials to criterion on the out-of-phase day in Experimental mice compared to Cre-negative controls. **d**, Top: experimental timeline during odor to texture shifts. mPFC and mIC PV INs are stimulated during each RS trial (in-phase one day, out-of-phase the other). Bottom: there is no significant differences in the trials needed to reach the learning criterion between days or groups. * p < 0.05, ** p < 0.01.

After five weeks, we tested mice on texture→odor shifts over two days (Figure 5c). The IA cue on the first day was randomly assigned. Mice received optogenetic excitation in the mIC and mPFC on both days. On either Day 1 or Day 2 (randomly assigned and counterbalanced across mice), mice received in-phase (IP) 40Hz stimulation during RS trials. On the other day, mice received 40Hz stimulation during RS trials that was out-of-phase (OOP) between the mIC and mPFC. Experimental (ChR2+) mice performed significantly worse during OOP stimulation compared to IP (p=0.0057), as measured by an increase in the number of trials to reach the learning criterion (8/10 trials correct). Experimental (ChR2+) mice performed significantly worse than (ChR2-negative) controls during OOP stimulation (p=0.028). There was no significant difference between control and experimental mice when they received IP stimulation (p=0.74), and there was no difference between controls on the IP and OOP stimulation days. Thus, artificially shifting mIC-mPFC PV IN gamma synchronization away from an in-phase pattern disrupts the learning of texture→odor shifts.

Next, we performed a similar experiment in a new cohort of mice, but delivered IP or OOP stimulation during odor→texture (rather than texture→odor) shifts (Figure 5d). In this case, performance was almost identical between control and experimental mice on both days, and between experimental mice when they received IP vs. OOP stimulation. Thus, the disruptive effect of OOP stimulation is specific for texture→odor shifts.

## DISCUSSION

The brain is organized as a network across many scales, and understanding the nature of these network interactions is crucial for understanding how the brain performs various computations and functions. Here, we elucidate how exactly the mIC and mPFC interact to generate flexible behavior. Our results suggest three key principles. First, mIC input to mPFC is not required for learning simple associations, but is specifically required for learning rule shifts. Second, this role is modality-specific (important for texture-to-odor, but not odor-to-texture shifts). Third, in-phase gamma synchrony between PV INs in the mIC and mPFC is an essential component of this interaction.

### Gamma synchrony and inter-regional communication

The role of gamma oscillations and their synchronization across brain regions has been an enduring controversy in systems neuroscience. Our prior work showed that gamma synchronization between PV INs in the left and right mPFC is necessary for learning rule shifts ^17^, but it was not clear if this extended to brain regions beyond the mPFC. Furthermore, in general, the concept that gamma-frequency neuronal activity can synchronize across widely separated brain regions has been challenging to demonstrate. Most demonstrations have involved synchronization between LFPs in different regions ^25,26^, or phase locking between neuronal spikes in one region and gamma-frequency components of LFPs believed to reflect activity in another region ^26,27^. Here, we not only identify gamma synchronization between voltage signals from specific neuronal populations located in two widely separated regions, but go on to show that this synchronization manifests in a behaviorally-specific manner, and confirm that it is causally necessary for behavior.

In-phase PV IN gamma synchrony between the mIC and mPFC increases during, and is specifically necessary for, texture→odor, but not odor→texture, shifts. This lines up with our observation that mIC-to-mPFC communication is necessary for texture→odor, but not odor→texture, shifts. Together, these observations suggest that this pattern of gamma synchrony is specifically required for mIC-to-mPFC communication. The idea that gamma synchrony is necessary for effective inter-regional communication is at the core of the ‘Communication Through Coherence (CTC)’ hypothesis ^21,22^. Thus, our findings are broadly consistent with the idea that mIC-mPFC PV IN gamma synchrony facilitates mIC-to-mPFC communication via CTC.

That said, the fact that texture→odor shifts are associated with an enrichment of mIC-mPFC PV IN phase differences between −45 and +45 degrees is somewhat surprising. Since these two structures are ∼3-4 mm apart, one might have expected this condition to be associated with an enrichment of phase differences of at least 5 msec (corresponding to 72 degrees) due to the axonal delay between mIC and mPFC. We offer two thoughts about this. First, pairs of bidirectionally coupled oscillators tend to synchronize either in-phase (0 degree phase lag) or anti-phase (+/− 180 degree phase lag), even when their coupling involves moderate phase delays. Of course, the mIC-mPFC circuit is much more complex than a simple pair of coupled oscillators. In this context, it would be interesting to study the mechanism of this near-zero-lag synchronization, for example, by testing how inhibiting connections in either the mIC-to-mPFC or mPFC-to-mIC direction affects patterns of gamma synchrony. We previously found that gamma synchronization between the two prefrontal hemispheres depends critically on callosal GABAergic projections from PV neurons ^18^, raising the possibility that similar, long-range GABAergic projections from PV neurons play a role here. Once the specific connections responsible for this pattern of synchronization have been identified, computational models might yield insights into how this in-phase pattern of synchronization emerges.

Second, in-phase gamma synchrony may not function simply to align the arrival of mIC signals with the phase of gamma associated with maximal mPFC excitability, as posited by CTC. This is because if PV IN activity aligns approximately in-phase between the mIC and mPFC, this likely means that the phase of peak excitability occurs at approximately the same time in both structures. Based on this, due to axonal delays, signals from the mIC would not arrive in the mPFC at the ‘optimal’ phase, corresponding to peak downstream excitability. Rather, in-phase synchronization may serve to align the arrival of signals from the mIC to gamma phases when specific computations are being performed or specific output signals are being generated in the mPFC. Future experiments which measure, for example, the timing of activity in mIC-mPFC projection neurons relative to gamma activity in the mPFC, might help clarify this issue.

Interestingly, OOP PV IN stimulation disrupted texture→odor shifts, but neither IP nor OOP stimulation has any effect on odor→texture shifts. This suggests that the predominately antiphase mode of mIC-mPFC PV IN gamma synchrony we observe during the pre-decision period of odor→texture shifts is not necessary for these shifts. This raises the question of whether this pattern of antiphase synchrony reflects a mode of communication that is not necessary for this behavior (possibly communication from mPFC to mIC), or whether it represents a form of active decoupling of these two regions.

Importantly, within the predecision period of correct trials during texture→odor shifts, gamma activity in PV INs in the mIC and mPFC is not synchronized in-phase the entire time. Rather, periods of in-phase mIC-mPFC PV IN gamma synchrony occur more often, and when they occur, the degree of synchronization (measured by the correlation value) is stronger. It is possible that these periods of strong in-phase synchronization correspond to some kind of network state during which specific types of information are transmitted from mIC to mPFC. This could be explored by recording activity of mIC-mPFC projection neurons simultaneous with measurements of gamma synchrony between these structures.

### The role of the prefrontal cortex in cognitive flexibility

Cognitive flexibility refers to the ability to alter behavioral strategies to adapt to changes in the environment. Cognitive flexibility is known to depend critically on the prefrontal cortex, and the prefrontal cortex is known to interact with numerous other regions, but the details of this have been unclear. Specifically, one possible model is that the prefrontal cortex itself stores and updates strategies to produce cognitive flexibility. Alternatively, the prefrontal cortex may simply monitor behavioral outcomes in order to generate output signals that direct other regions to learn new strategies and/or unlearn outdated ones. Yet another possibility is that the prefrontal cortex primarily acts to suppress outdated strategies, which effectively allows new ones to emerge.

Some of the preceding functions require the prefrontal cortex to receive and utilize input related to the specific cues involved in the behavioral strategies, while others do not. Here we show that input from the mIC contributes to cognitive flexibility during texture to odor, but not odor to textue, shifts. This suggests that the prefrontal cortex is not agnostic to the nature of the cues involved in strategies, i.e., the prefrontal cortex is not simply monitoring behavioral outcomes and creating output signals that support flexible behavior in a manner that is independent of cue information. Rather, the details of the behavioral strategy, e.g., the particular cues that it depends on, seem to be actively transmitted to the prefrontal cortex and used to promote flexibility. This is consistent with models in which the prefrontal cortex actively represents associations between specific cues and reward, and/or guides choice selection based on specific cues, as opposed to simply monitoring the overall rate of rewards and representing levels of uncertainty, etc.

## Conclusion

Communication and coordination between regions is an essential component of brain function. Here we show that in-phase gamma-frequency synchronization plays a key role in communication between the insula and prefrontal cortex that is necessary when animals learn new rules involving specific modalities. This reveals a key principle underlying inter-regional coordination, and also provides details about exactly how sensory cue-related information is transmitted to higher-order brain regions in order to contribute to cognitive processes. Future work could explore both the underlying mechanism and neural information processing function of this pattern of temporal coordination.

**Table 1:** Summary of statistics for Figures 1-3.

| Panel | Details |  | Mixed Effects |  |  | Multiple Comparisons |  |
| --- | --- | --- | --- | --- | --- | --- | --- |
|  | Days of Rule Shifting | Optogenetic Inhibition | Fixed effects (type III) | P value | F (DFn, DFd) | Comparison | Adjusted P Value |
| Figure 1d | 3 | Day 3 durring T→O RS | Day | 0.0004 | F (1.543, 12.35) = 18.15 | Day 1 |  |
|  |  |  | Group | 0.0322 | F (1.000, 8.000) = 6.701 | Experimental (eNpHR+) vs. Control (eYFP+) | >0.9999 |
|  |  |  | Day x Group | 0.0092 | F (1.726, 13.81) = 7.135 |  |  |
|  |  |  |  |  |  | Day 2 |  |
|  |  |  |  |  |  | Experimental (eNpHR+) vs. Control (eYFP+) | >0.9999 |
|  |  |  |  |  |  | Day 3: Inhibition |  |
|  |  |  |  |  |  | Experimental (eNpHR+) vs. Control (eYFP+) | 0.0001 |
|  |  |  |  |  |  | Experimental (eNpHR+) |  |
|  |  |  |  |  |  | Day 1 vs. Day 2 | 0.8147 |
|  |  |  |  |  |  | Day 1 vs. Day 3: Inhibition | <0.0001 |
|  |  |  |  |  |  | Day 2 vs. Day 3: Inhibition | <0.0001 |
|  |  |  |  |  |  | Control (eYFP+) |  |
|  |  |  |  |  |  | Day 1 vs. Day 2 | 0.8147 |
|  |  |  |  |  |  | Day 1 vs. Day 3: Inhibition | 0.11 |
|  |  |  |  |  |  | Day 2 vs. Day 3: Inhibition | 0.0293 |
| Figure 2a | 3 | Day 3 durring O→T RS | Day | 0.1774 | F (2, 10) = 2.066 | Day 1 |  |
|  |  |  | Group | 0.1375 | F (1, 5) = 3.122 | Experimental (eNpHR+) vs. Control (eYFP+) | 0.3944 |
|  |  |  | Day x Group | 0.7136 | F (2, 7) = 0.3542 |  |  |
|  |  |  |  |  |  | Day 2 |  |
|  |  |  |  |  |  | Experimental (eNpHR+) vs. Control (eYFP+) | 0.1204 |
|  |  |  |  |  |  | Day 3: Inhibition |  |
|  |  |  |  |  |  | Experimental (eNpHR+) vs. Control (eYFP+) | 0.6226 |
|  |  |  |  |  |  | Experimental (eNpHR+) |  |
|  |  |  |  |  |  | Day 1 vs. Day 2 | 0.6459 |
|  |  |  |  |  |  | Day 1 vs. Day 3: Inhibition | 0.4332 |
|  |  |  |  |  |  | Day 2 vs. Day 3: Inhibition | 0.9306 |
|  |  |  |  |  |  | Control (eYFP+) |  |
|  |  |  |  |  |  | Day 1 vs. Day 2 | 0.1948 |
|  |  |  |  |  |  | Day 1 vs. Day 3: Inhibition | 0.5931 |
|  |  |  |  |  |  | Day 2 vs. Day 3: Inhibition | 0.6932 |
| Figure 2b | 2 | Day 2 durring Odor IA | Day | 0.0149 | F (1, 18) = 7.244 | Day 1 |  |
|  |  |  | Group | 0.2198 | F (1, 18) = 1.616 | Experimental (eNpHR+) vs. Control (eYFP+) | 0.3397 |
|  |  |  | Day x Group | 0.9093 | F (1, 18) = 0.01336 |  |  |
|  |  |  |  |  |  | Day 2: Inhibition |  |
|  |  |  |  |  |  | Experimental (eNpHR+) vs. Control (eYFP+) | 0.4245 |
|  |  |  |  |  |  | Experimental (eNpHR+) |  |
|  |  |  |  |  |  | Day 1 vs. Day 2: Inhibition | 0.0519 |
|  |  |  |  |  |  | Control (eYFP) |  |
|  |  |  |  |  |  | Day 1 vs. Day 2: Inhibition | 0.0982 |
| Figure 3c | 3 | Day 3 durring T→O RS | Day | 0.0063 | F (2, 12) = 7.968 | Day 1 |  |
|  |  |  | Group | 0.1187 | F (1, 6) = 3.310 | Experimental (eNpHR+) vs. Control (eYFP+) | 0.7075 |
|  |  |  | Day x Group | 0.1041 | F (2, 9) = 2.939 |  |  |
|  |  |  |  |  |  | Day 2 |  |
|  |  |  |  |  |  | Experimental (eNpHR+) vs. Control (eYFP+) | 0.9526 |
|  |  |  |  |  |  | Day 3: Inhibition |  |
|  |  |  |  |  |  | Experimental (eNpHR+) vs. Control (eYFP+) | 0.0086 |
|  |  |  |  |  |  | Experimental (eNpHR+) |  |
|  |  |  |  |  |  | Day 1 vs. Day 2 | 0.9784 |
|  |  |  |  |  |  | Day 1 vs. Day 3: Inhibition | 0.0019 |
|  |  |  |  |  |  | Day 2 vs. Day 3: Inhibition | 0.0012 |
|  |  |  |  |  |  | Control (eYFP+) |  |
|  |  |  |  |  |  | Day 1 vs. Day 2 | 0.9887 |
|  |  |  |  |  |  | Day 1 vs. Day 3: Inhibition | 0.5827 |
|  |  |  |  |  |  | Day 2 vs. Day 3: Inhibition | 0.6705 |

**Table 2:** Summary of statistics for Figure 4.

| Panel | Time Period | Statistical test |  |  | Multiple comparison |  |
| --- | --- | --- | --- | --- | --- | --- |
|  |  |  |  |  | Comparison | Adjusted P-value |
| Figure 4e | Pre-Decsion | Mixed Effect |  |  |  |  |
|  |  | Fixed effects (type III) | P value | F (DFn, DFd) | Correct Trials |  |
|  |  | Trial Type | 0.8788 | F (1, 16) = 0.02399 | Texture to Odor vs. Odor to Texture | 0.1823 |
|  |  | Shift Type | 0.2077 | F (1, 16) = 1.724 |  |  |
|  |  | Trial Type x Shift Type | 0.0047 | F (1, 16) = 10.79 | Error Trials |  |
|  |  |  |  |  | Texture to Odor vs. Odor to Texture | 0.005 |
|  |  |  |  |  | Texture to Odor |  |
|  |  |  |  |  | Correct Trials vs. Error Trials | 0.0271 |
|  |  |  |  |  | Odor to Texture |  |
|  |  |  |  |  | Correct Trials vs. Error Trials | 0.0418 |
| Figure 4g | Pre-Decsion Correct Trials | Chi-square |  |  | NA | NA |
|  |  | Comparison | Sides | P value |  |  |
|  |  | In-Phase [-45, 45] vs Anti-Phase [-135,-225] vs Anti-Phase [135,225] | Two | 0.0014 |  |  |
| Figure 4h | Pre-Decsion Correct Trials | 2-Way ANOVA |  |  | NA | NA |
|  |  | Source of Variation | P value | F (DFn, DFd) |  |  |
|  |  | Interaction | <0.0001 | F (32, 264) = 3.435 |  |  |
|  |  | Phase Shift | <0.0001 | F (32, 264) = 7.696 |  |  |
|  |  | Behavior Shift | 0.6296 | F (1, 264) = 0.2331 |  |  |
| Figure 4i | Pre-Decsion Error Trials | 2-Way ANOVA |  |  | NA | NA |
|  |  | Source of Variation | P value | F (DFn, DFd) |  |  |
|  |  | Interaction | 0.9864 | F (32, 264) = 0.5177 |  |  |
|  |  | Phase Shift | <0.0001 | F (32, 264) = 5.861 |  |  |
|  |  | Behavior Shift | 0.6962 | F (1, 264) = 0.1527 |  |  |
| Figure 4j | Post Outcome | Mixed Effect |  |  |  |  |
|  |  | Fixed effects (type III) | P value | F (DFn, DFd) | Correct Trials |  |
|  |  | Trial Type | 0.0028 | F (1, 4) = 43.19 | Texture to Odor vs. Odor to Texture | 0.202 |
|  |  | Shift Type | 0.3796 | F (1, 4) = 0.9736 |  |  |
|  |  | Trial Type x Shift Type | 0.383 | F (1, 4) = 0.9584 | Error Trials |  |
|  |  |  |  |  | Texture to Odor vs. Odor to Texture | 0.9958 |
|  |  |  |  |  | Texture to Odor |  |
|  |  |  |  |  | Correct Trials vs. Error Trials | 0.0004 |
|  |  |  |  |  | Odor to Texture |  |
|  |  |  |  |  | Correct Trials vs. Error Trials | 0.0014 |
| Figure 4k | Post Outcome Correct Trials | 2-Way ANOVA |  |  | NA | NA |
|  |  | Source of Variation | P value | F (DFn, DFd) |  |  |
|  |  | Interaction | 0.9988 | F (32, 264) = 0.3955 |  |  |
|  |  | Phase Shift | <0.0001 | F (32, 264) = 21.70 |  |  |
|  |  | Behavior Shift | 0.4199 | F (1, 264) = 0.6527 |  |  |
| Figure 4k | Post Outcome Error Trials | 2-Way ANOVA |  |  | NA | NA |
|  |  | Source of Variation | P value | F (DFn, DFd) |  |  |
|  |  | Interaction | 0.7939 | F (32, 264) = 0.7837 |  |  |
|  |  | Phase Shift | <0.0001 | F (32, 264) = 38.39 |  |  |
|  |  | Behavior Shift | 0.4427 | F (1, 264) = 0.5910 |  |  |
| Figure 4k | ITI Correct Trials | 2-Way ANOVA |  |  | NA | NA |
|  |  | Source of Variation | P value | F (DFn, DFd) |  |  |
|  |  | Interaction | 0.9578 | F (32, 264) = 0.6011 |  |  |
|  |  | Phase Shift | <0.0001 | F (32, 264) = 33.30 |  |  |
|  |  | Behavior Shift | 0.6252 | F (1, 264) = 0.2392 |  |  |
| Figure 4k | ITI Error Trials | 2-Way ANOVA |  |  | NA | NA |
|  |  | Source of Variation | P value | F (DFn, DFd) |  |  |
|  |  | Interaction | 0.6318 | F (32, 264) = 0.8964 |  |  |
|  |  | Phase Shift | <0.0001 | F (32, 264) = 31.29 |  |  |
|  |  | Behavior Shift | 0.6945 | F (1, 264) = 0.1546 |  |  |

**Table 3:** Summary of statistics for Figure 5.

| Panel | Details |  | 2-Way ANOVA |  |  | Multiple Comparisons |  |
| --- | --- | --- | --- | --- | --- | --- | --- |
|  | Days Rule-Shifting | Optogenetic Excitation | Source of Variation | P value | F (DFn, DFd) | Comparison | Adjusted P Value |
| Figure 5c | 2 | In phase or Out of Phase during T→O | *Interaction* | 0.168 | F (1, 20) = 2.047 | *In-Phase* |  |
|  |  |  | *Excitation Type* | 0.0443 | F (1, 20) = 4.606 | *Experimental (ChR2+) vs. Control (ChR2-)* | 0.7395 |
|  |  |  | *Group* | 0.0709 | F (1, 20) = 3.639 |  |  |
|  |  |  |  |  |  | *Out-of-phase* |  |
|  |  |  |  |  |  | *Experimental (ChR2+) vs. Control (ChR2-)* | 0.0285 |
|  |  |  |  |  |  | *Experimental (ChR2+)* |  |
|  |  |  |  |  |  | *In-Phase vs. Out-of-phase* | 0.0057 |
|  |  |  |  |  |  | *Control (ChR2-)* |  |
|  |  |  |  |  |  | *In-Phase vs. Out-of-phase* | 0.666 |
| Figure 5d | 2 | In phase or Out of Phase during O→T | *Interaction* | 0.5894 | F (1, 20) = 0.3008 | *In-Phase* |  |
|  |  |  | *Excitation Type* | 0.1155 | F (1, 20) = 2.708 | *Experimental (ChR2+) vs. Control (ChR2-)* | 0.2289 |
|  |  |  | *Group* | 0.2416 | F (1, 20) = 1.456 |  |  |
|  |  |  |  |  |  | *Out-of-phase* |  |
|  |  |  |  |  |  | *Experimental (ChR2+) vs. Control (ChR2-)* | 0.6467 |
|  |  |  |  |  |  | *Experimental (ChR2+)* |  |
|  |  |  |  |  |  | *In-Phase vs. Out-of-phase* | 0.3534 |
|  |  |  |  |  |  | *Control (ChR2-)* |  |
|  |  |  |  |  |  | *In-Phase vs. Out-of-phase* | 0.1941 |

## ACKNOWLEDGEMENTS

We acknowledge technical advice and assistance from Adam Jackson, Ruchi Malik, Xiyu Zhu, Caitriona Costello and Christine Liu. This work was supported by NIH grants R01NS116594 (to V.S.S.), R01MH121342 (to V.S.S.), and R01MH129835 (to V.S.S.).

## DECLARATION OF INTERESTS

The authors declare no competing interests.

## METHODS

### Mice

NIH guidelines and a protocol approved by the Institutional Animal Care & Use Committee (IACUC) at the University of California, San Francisco were followed for all animal care, procedures and experiments. Mice were group housed (2-5 siblings) with full access to food and water in a normal 12hr light-dark cycle room kept at 22-24°C until Rule-Shifting experiments began. All experiments used wild-type, PV-flp, and PV-flp/Ai14 lines on a C57Bl/6 obtained from Jackson Laboratories. Adult male and female mice were used, and were 8-12 weeks at the time of surgery and 16-20 weeks old during behavioral experiments.

### Cloning of Viral Constructs

Viruses pAAV-EF1a-Cre, AAV5-Ef1a-DIO-eNPHR-eYFP, and AAV5-Ef1a-DIO-eYFP were all purchased from Addgene. AAV2-Syn-tdTomato (1.23E+12 vg/mL) were purchased from SignaGen Laboratories. We received AAV1-CAG-fDIO-Ace2N-4AA-mNeon (2.23E+13 vg/mL) from Mark J. Schnitzer (Stanford University) or the pAAV-CAG-fDIO-Ace2N-4AA-mNeon plasmid which was then packaged with stereotype AAV1 by Virovek (Houston, TX).

### Surgery

All mice were anaesthetized using isoflurane (induction: 2.5% / maintenance: 1.2 - 1.5%; in 95% oxygen) then placed on a stereotaxic frame (David Kopf Instruments) with a heating pad to maintain body temperature. The mouse’s head was shaved with a electric razor and sterilized with ethanol and betadine. Lidocaine (1:4 dilution in saline) was injected underneath the scalp, and meloxicam (1:10 dilution) and Ethiqa XR were injected subcutaneously for analgesia. An incision was made to expose the skull. The scalp was pushed to the sides and periosteum was removed from the dorsal skull surface. Necessary regions of the skull were drilled to create an opening for viral implants and/or injections. Using a micro-syringe pump (World Precision Instruments, UMP3 UltraMicroPump), viruses were infused at 100nL min through a 35-gauge, beveled injection needle (World Precision Instruments). The needle was kept at the injection site for 3-5min post injection before being withdrawn. The skull was recleaned with saline. Implants were placed one at a time at targeted sites, and affixed to the skull using Metabond Quick Adhesive Cement (Parkell).

#### mIC-to-mPFC cell body inhibition

For optogenetic inhibition of mIC-to-mPFC cell bodies, wild-type C57 mice were injected bilaterally in mPFC prelimbic and infralimbic cortices (1.7 AP; ±0.3 ML; −2.0, −2.25, −2.5 DV mm) and mIC (0.86 AP; ±3.4 ML; −4.0 DV mm) (all coordinates relative to bregma). In each hemisphere, the mPFC was injected with 3×0.2µL of the retrograde virus AAVrg-EF1a-Cre (Addgene). Using a lateral facing beveled needle, the mIC was injected in each hemisphere with 600µL of AAV5-Ef1a-DIO-eNPHR-eYFP or AAV5-Ef1a-DIO-eYFP (Addgene). Implants with a 200/240µm (core/outer) diameter, NA=0.22 (Doric Lenses, MFC_200/240-0.22_2.8mm_ZF1.25(G)_FLT), were placed bilaterally in the mIC (0.86 AP; ±3.35 ML; −3.75 DV mm). To allow sufficient time for viral expression, all experiments began 8-10 weeks after surgery.

#### mIC to mPFC Terminal Inhibition

For mIC to mPFC terminal optogenetic inhibition experiments, wild-type C57 mice were injected bilaterally in the prelimbic and infralimbic regions of mPFC (1.7 AP; ±0.3 ML; −2.0, −2.25, −2.5 DV mm) and mIC (0.86 AP; ±3.4 ML; −4.0 DV mm) relative to bregma. In each hemisphere, we injected mPFC with 3×0.2µL of the retrograde virus AAVrg-EF1a-Cre (Addgene). Using a lateral facing beveled needle, we injected mIC in each hemisphere with 600µL of AAV5-Ef1a-DIO-eNPHR-eYFP or AAV5-Ef1a-DIO-eYFP (Addgene). Implants with a 200/240µm (core/outer) diameter, NA=0.22 (Doric Lenses, MFC_200/240-0.22_2.8mm_ZF1.25(G)_FLT), were placed bilaterally in the mPFC at a ±12° angle (1.7 AP, ±0.76, −2.14 DV). To allow time for viral expression, experiments began 8-10 weeks after surgery.

#### TEMPO recording in mIC and mPFC

For TEMPO experiments, PV-Flp mice were injected unilaterally in the left mPFC prelimbic and infralimbic cortices (1.7 AP; −0.3 ML; −2.0, −2.25, −2.5 DV mm) and left mIC (0.86 AP; −3.4 ML; −4.0 DV mm) (all coordinates relative to bregma) with a 2:1 mixture of virus to drive GEVI expression (AAV1-CAG-fDIO-Ace2N-4AA-mNeon; Schnitzer Lab) and AAV2-Syn-tdTomato (1.23E+ 12 vg/mL; SignaGen Laboratories). mPFC was injected with 3×0.2µL of this mixture. Using a lateral facing beveled needle, we injected mIC with 500µL of the same mixture. Implants with a 400/430µm (core/outer) diameter, NA=0.48 (Doric Lenses, MFC_400/430-0.48_2.8mm_ZF1.25_FLT), were implanted in the left mPFC (1.7 AP; ±0.3 ML; −2.0 DV mm) and left mIC (0.86 AP; ±3.35 ML; −3.75 DV mm). To allow time for viral expression, experiments began about 5 weeks after surgery.

#### mIC and mPFC In-Phase and Out-of-Phase optogenetic stimulation

For mIC and mPFC PV IN optogenetic stimulation experiments, PV-Cre^+/−^(experimental) and wild-type C57 (control) mice were injected bilaterally in the prelimbic and infralimbic cortices (1.7 AP; ±0.3 ML; −2.0, −2.25, −2.5 DV mm) and mIC (0.86 AP; ±3.4 ML; −4.0 DV mm) (all coordinates relative to bregma). In each hemisphere, we injected mPFC with 3×0.2µL of AAV5-EF1a-DIO-ChR2-eYFP. Using a lateral facing beveled needle, we injected mIC in each hemisphere with 600µL of AAV5-EF1a-DIO-ChR2-eYFP. Dual fiber optic implants with a 200/240µm (core/outer) diameter, NA=0.22 (Doric Lenses, DFC_200/240-0.22_2.3mm_GS 0.7_FLT), were placed bilaterally in the mPFC. Fiber optic implants with a 200/240µm (core/outer) diameter, NA=0.22 (Doric Lenses, MFC_200/240-0.22_2.8mm_ZF1.25(G)_FLT), were placed bilaterally in the mIC (−0.21 AP; ±3.4 ML; −3.9 DV mm) at a 16° angle. To allow time for viral expression, experiments began 4-5 weeks after surgery.

### Behavior

#### Rule Shifting Task

A detailed description of this task can be found in Cho et al., 2015. In this task, mice learn a rule or ‘Initial Association’ (IA) between a texture or odor cue and the location of a food reward, then switch to a new rule (‘Rule Shift’; RS) based on a cue from the opposing modality. In all experiments, the experimenter was blind to genotype and/or virus injected while running mice on this task.

Mice were placed with litter mates in a reverse light/dark cycle for 24-48 hours with normal food and water. Full diet 20mg food pellets (Dustless Precision Pellets, 20mg, Rodent Purified diet; Bio-Serv) were sprinkled into their cages during this time to acclimate them to the food reward, prior to beginning food deprivation.

We mixed ground dried spices (McCormick; garlic or coriander) representing the odor cues into the digging media (sand or litter) (odor concentration: ∼0.1% by volume). We also mixed ground food pellet powder into this mixture (∼0.01% of the volume) to prevent mice from being able to smell the hidden food pellet.

During the period of food deprivation, mice were singly housed under a reverse light cycle for approximately one week. Food consumption was restricted to maintain mice at 80-85% of their pre-restriction weight. Mice were specifically given pellets, placed in the middle of the media within two bowls, one filled with sand (Mosser Lee White Sand Soil Cover) plus an odor of garlic or coriander, the other with litter (Natural Integrity Clumping Clay cat litter) with the other odor.

After 2-3 days of food deprivation, mice generally reach their target weight. They are then run through a single day ‘habituation’ session. During habituation, mice run 10 consecutive trials with baited food bowls. In these trials, the mouse finds a reward while digging in all possible media combinations and each location (left or right) at least once. Mice are free to sample both bowls as they please until they find/eat the reward and are then placed back into the holding cage. If mice fail to dig in the baited bowl, they are considered to have failed habituation, and the habituation protocol is repeated the following day; if they fail again they are removed from the experiment.

On the following 2-3 days mice run the task. To determine which odor/texture combination and side (left or right) baited bowls will be placed on each trial, we used a custom MATLAB script that randomly assigns these values, with the requirement that the same odor/texture combination and spatial location of reward (left or right side) does not repeat on more than 2 consecutive trials.

At the start of each trial, the mouse is moved from the holding cage to its home cage which contains two bowls, only one of which is baited with a hidden food reward; the mouse digs in a bowl and is then placed back into the holding cage. A trial is considered correct if the mouse digs in the baited bowl. The intertrial intervals following correct trials are about 10s, allowing time for the bowls to be replaced. A trial is considered incorrect when a mouse digs in the non-baited bowl; in this case, the baited bowl is removed and the mouse is free to dig in the non-baited bowl until it loses interest; at that point the mouse is moved to the holding cage for a 40s intertrial interval (‘time out’).

On each day mice must first learn an initial association (IA) between one cue and the location of the hidden food reward. Once mice get 8/10 consecutive trials correct, they are considered to have learned the rule. If on a non-manipulation day, mice fail to learn the IA within 15 trials, they fail that day and are run again the next day; if they fail a second time they are removed from the experiment. Once IA is learned, the experimenter moves on to the rule-shift (RS); here the rule is shifted to the opposing modality (example: If the IA is based on an odor cue then the RS uses a texture cue, and vice versa). Once mice get 8/10 consecutive trials correct, they are considered to have learned the rule-shift. Under normal conditions, mice typically learn the IA in about 10 trials and the RS in about 10-15 trials.

For analyses of neural activity, we considered the onset of digging as the time of the decision.

#### Optogenetic Inhibition

Continuous light for eNpHR stimulation was delivered during trials using a 532nm green laser coupled to two mono fiber-optic cannulas (Doric Lenses, Inc) through a 200µm diameter Splitter Branching Fiber optic patch cord with mating sleeves (Doric Lenses, Inc.). The final light power was adjusted to ∼2.5mW for each branch (∼5mW total across both branches). While the mouse was in the holding cage, before the start of a trial, the laser was manually turned on, and the mouse was placed in its home cage for the start of the trial. Once a mouse was placed back in the holding cage the laser was turned off; the laser remained off for the remainder of the intertrial interval, including time-outs.

#### Optogenetic Excitation (In-Phase vs. Out-of-Phase)

A single 40Hz train of 5ms pulses was delivered to both fibers to produce in-phase stimulation. Out-of-phase stimulation was delivered using a 10ms (144 degree) offset between trains in the two fibers. ChR2 stimulation during trials was delivered via a 473nm blue laser coupled with two mono fiber-optic cannulas (Doric Lenses, Inc) through a 200µm diameter Splitter Branching Fiber optic patch cord with mating sleeves (Doric Lenses, Inc.). For optogenetic excitation, we adjusted the laser to deliver a continuous (non-pulsed) light power ∼1.3mW for each fiber (∼5.2mW total across the four fibers), then switched to 40Hz pulsing. While the mouse was in the holding cage, before the start of each trial, the laser was manually turned on, and the mouse was placed in its home cage for the start of the trial. Once a mouse was placed back in the holding cage the laser was turned off; the laser remained off during any portion of the intertrial intervals, including time-outs.

### Histology and Imaging

All mice used for experiments were anesthetized with Euthasol, then transcardinally perfused with 0.01M PBS; once the blood ran clear, mice were perfused with 4% paraformaldehyde (PFA) in PBS until muscle spasms stopped or the body was stiff. Brains were dissected and stored in 4% PFA overnight at 4°C, then rinsed with PBS and transferred to 30% sucrose solution made in 0.01M PBS. We waited at least 1-2 days (for brains to sink in the solution) before slicing. Brains were sliced at 60µm on a Leica SM2000 Microtome and mounted on slides with Fluoromount G with DAPI (Southern Biotech, Birmingham, AL). All imaging was performed on a BZ-X Series all-in-one fluorescence microscope (Keyance). We verified all mice included here had optical fibers and/or viral expression located in the proper regions.

### Transmembrane Electrical Measurements Performed Optically (TEMPO)

TEMPO was based on techniques originally described in Marshall et al., 2016, then modified by our lab as described in Cho et al., 2020, Cho et al., 2023, Jackson et al., 2024, Phensy et al., 2026 and detailed below.

#### Optical Apparatus

Fiber optic implants with a 400/430µm (core/outer) diameter, NA=0.48 (Doric Lenses, MFC_400/430-0.48_2.8mm_ZF1.25_FLT), were sterotaxically placed in the targeted region as described earlier. During recording, a mouse was plugged in using a fiber-optic patch cord (Doric Lenses, MFP_400/430/1100–0.48_2m_FC-ZF1.25) which was secured with a zirconia sleeve (Doric Lenses). This provided the light path between the mouse and a miniature, permanently-aligned optical bench, or ‘mini-cube’ (Doric Lenses, FMC5_E1(460–490)_F1(500–540)_E2(555–570)_F2(580–680)_S). Each fiber delivered excitation light and received emitted fluorescence from a recording site, and each recording site had its own implant, patch cord, mini-cubes, LEDs, photoreceivers and lock-in amplifiers.

Each mini-cube monitors two spectrally separate fluorophores. It does so using dichroic mirrors and clean up filters that match the excitation and emission spectra of the voltage sensor (Ace-mNeon: Excitation 460–490 nm, Emission 500–540 nm) and reference fluorophore (tdTomato: Excitation 555–570 nm, Emission 580–680 nm). Mini-cubes were permanently aligned and sealed. Each cube has 5 ports; one sample port that plugs into the animal, and two excitation and two emission lines. Each port has matching coupling optics and FC connectors.

The mini-cube was connected to fiber-coupled LEDs (Center wavelengths 490 nm and 565 nm, Thorlabs M490F3 and M565F3) via a patch cord (200 μm, NA=0.39; Thorlabs M75L01). LEDs were controlled using a 4-channel, 10kHz-bandwidth current source (Thorlabs DC4104), and the LED light intensity was adjusted to be ∼200µW (Ace-mNeon) and ∼100µW (tdTomato).

Both emission ports on the mini-cube were attached to an adjustable-gain photoreceiver (Femto, Berlin, Germany, OE-200-Si-FC; Bandwidth set to 7kHz, AC-coupled, ‘Low’ gain of ∼5×10^7 V/W) using a large-care high-NA fiber was used (600μm core, NA=0.48 (Doric lenses, MFP_600/630/LWMJ-0.48_0.5m_FC-FC)).

#### Modulation and lock-in detection

Each recording site had two LEDs. All four LEDs were sinusoidally modulated at distinct frequencies to reduce crosstalk between fluorophores with overlapping spectra. Lock-in amplifiers were used to demodulate corresponding photoreceiver outputs and remove noise using low pass filters. Each LED was driven using a modulated waveform, and we demodulated emitted fluorescence signals at the level of the photoreceiver output using the corresponding carrier frequency. This was done using a single lock-in amplifier (Stanford Research Systems, SR860). We used a different carrier frequency for each site and flurophore (mNeon site 1: 2 kHz; mNeon site 2: 2.5 kHz; tdTomato site 1: 3.5 kHz; tdTomato site 2: 4 kHz).

#### TEMPO recording

A multichannel real-time signal processor (Tucker-Davis Technologies, Alachua, Florida; RX-8) was used for digitization. This signal processor was controlled using the Synapse software package (Tucker-Davis Technologies) running on a Windows PC. Synapse saved data to files on the PC and synchronized recordings with simultaneously acquired video from a USB webcam (Ailipu Technology, Shenzhen, China, ELP-USB100W05MT-DL36).

#### TEMPO Analysis

Methods for TEMPO analysis are described in previous work (Phensy, A.J. et al) but are briefly described here.

Videos were scored to identify timepoints corresponding to the trial start, dig start, outcome, and trial end. The pre-decision period was defined as 1-3s seconds before dig start; the post decision period was defined as the time between the outcome and trial end.

Cross hemispheric gamma synchrony between PV INs was quantified using 250ms windows. First, following data acquisition, data was converted into a MATLAB structure using the SEV2Mat script from Tucker-Davis Technologies. Second, all signals were extracted: 1) left mIC mNeon, 2) left mIC tdTomato, 3) left mPFC mNeon 4) left mPFC tdTomato. Third, linear regression between broadband filtered (1.6 - 78.4 Hz) mNeon and tdTomato signals (recorded from the same site) was performed, to identify the shared noise component. This shared noise component was subtracted from each mNeon signal to ‘clean’ that signal; this allowed for non-voltage-related shared noise to be removed. Fourth, signals were band-pass filtered specifically within the gamma frequency band (30-50Hz) and an additional linear regression between mNeon and tdTomato was conducted to remove any additional noise. Lastly, we calculated the cross correlation between these cleaned filtered mNeon signals from mIC and mPFC across the entire session in 250ms windows to quantify gamma synchrony. We identified when the peak of the cross-correlation was between −45 to 45 degrees and used these values to quantify ‘zero-phase lag’ mIC-mPFC PV IN gamma synchrony. Comparisons were based on these cross-correlation values.

Using these values, group comparisons were made using a mixed effects model or paired nonparametric two tailed t-test. We corrected for multiple comparisons using statistical hypothesis testing (Tukey); if this correction was not possible due to 3 or less data points, each comparison stood on its own using a Fishers LSD test.

### General Data Statistics for Behavior

Graphpad Prism 10 was used to conduct statistical analysis (see figure legends for details). Quantitative data are shown as the mean; error bars are used to represent the standard error of the mean (s.e.m.). Measurements were taken from distinct samples (*P* = *, < 0.05; **, <0.01; ***, < 0.001; ****, < 0.0001); no asterisk implied no significant difference (*P* > 0.05). Sample sizes were based on number of mice available. Data distribution was assumed to be non-normal.

Group comparisons were made using mixed effects models (behavioral effects of optogenetic inhibition, levels of synchrony), 2-way-ANOVAs (behavioral effects of optogenetic excitation), or chi-square tests (fractions of timepoints with different phase lags). If an interaction and/or a main effect was present, we corrected for multiple comparisons using statistical hypothesis testing (Tukey); if this correction was not possible due to limited data points, each comparison stood on its own using a Fishers LSD test.

## Supplementary Figures

**Supplementary Figure 1:**
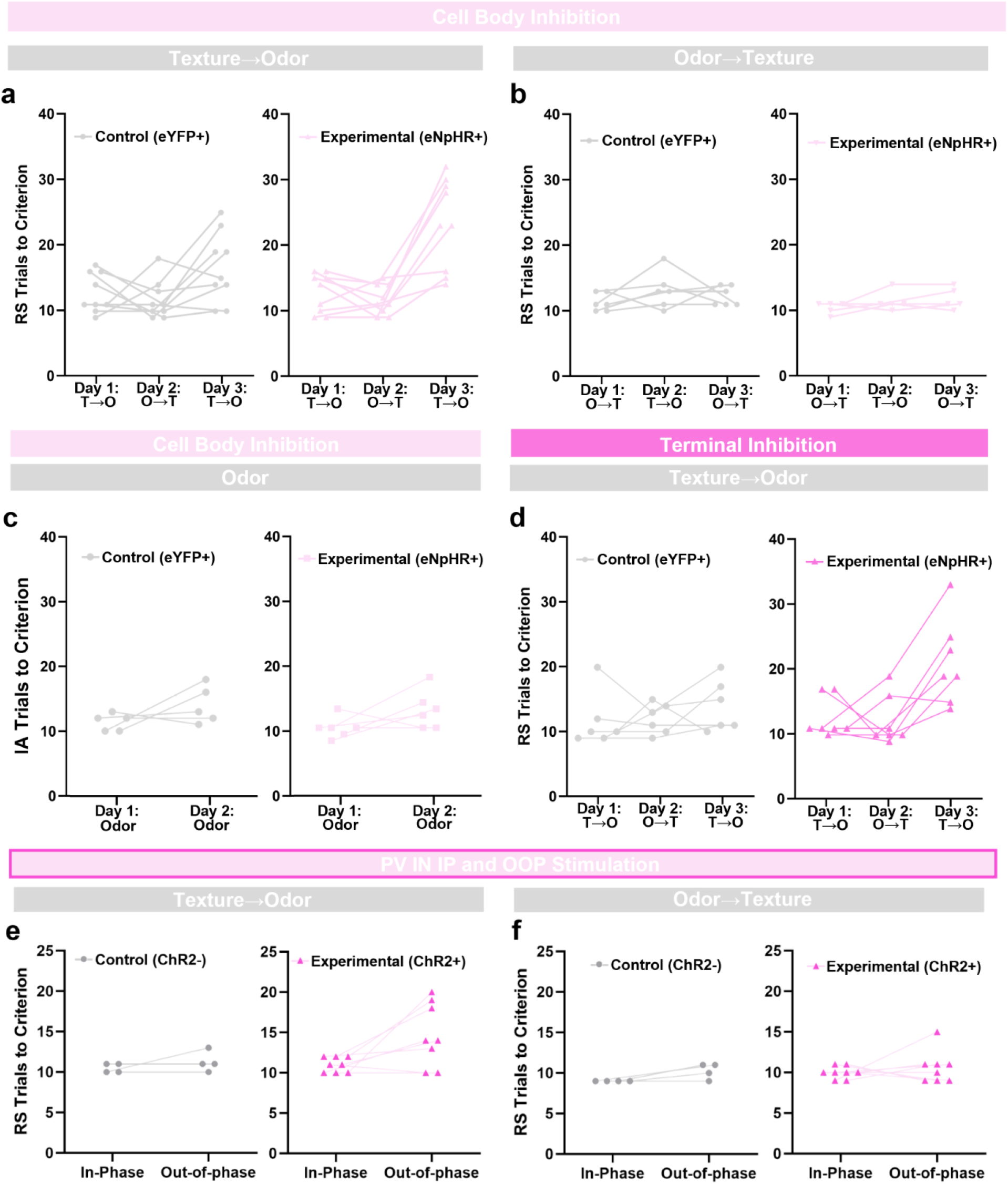
Behavior of individual mice during optogenetic manipulations. **a,** Behavior of mice receiving mIC to mPFC cell body inhibition while shifting from texture → odor on Day 3. **b,** Behavior of mice receiving mIC to mPFC cell body inhibition while shifting from odor → texture on Day 3. **c,** Behavior of mice receiving cell body inhibition when learning odor-based IA on Day 2. **d,** Behavior of mice receiving mIC to mPFC terminal inhibition while shifting from texture → odor. **e,** Behavior of mice when shifting from texture → odor during IP and OOP excitation of PV INs in mIC and mPFC. **f,** Behavior of mice when shifting from odor → texture during IP and OOP excitation of PV INs in mIC and mPFC.

**Supplementary Figure 2:**
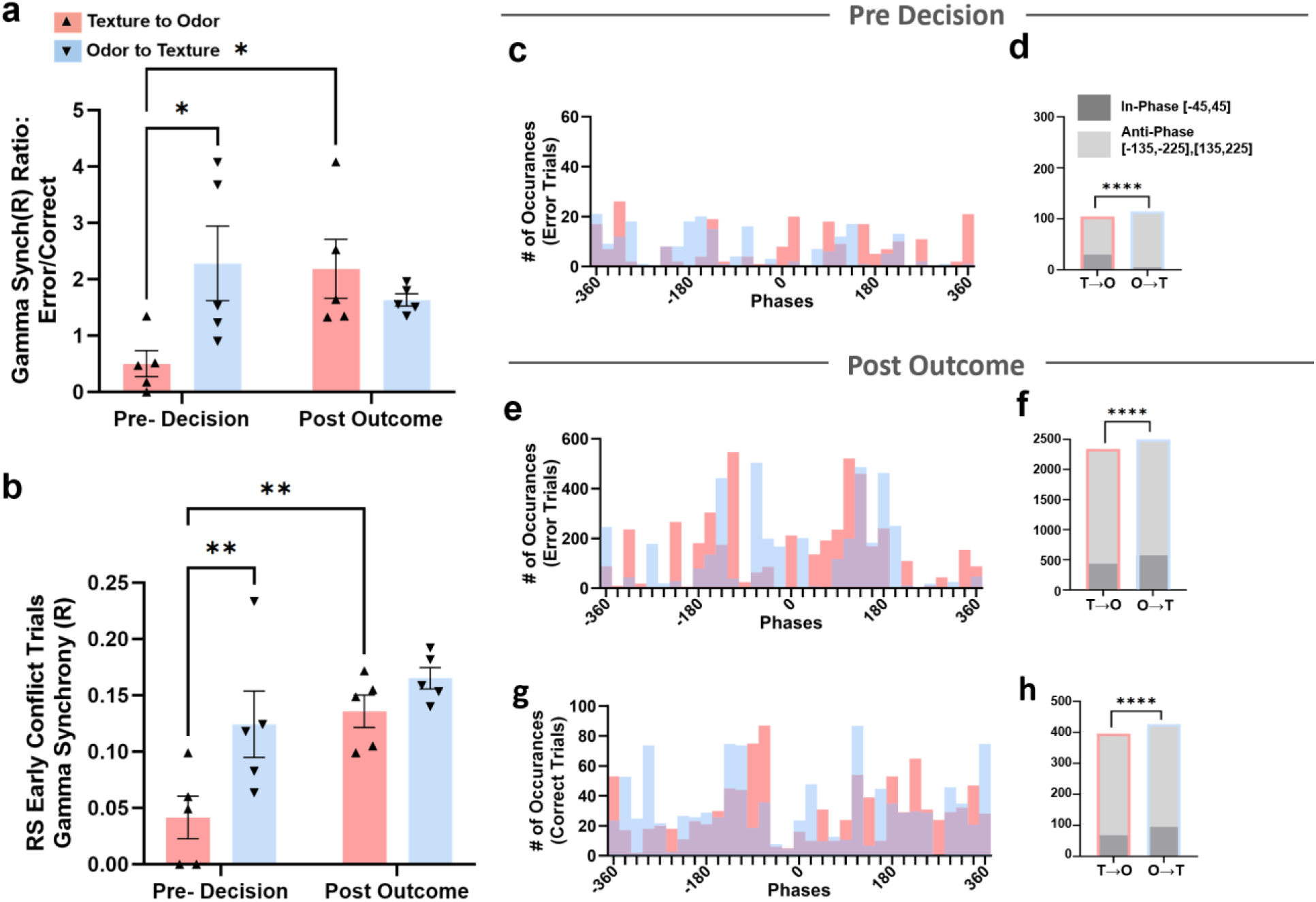
mIC-mPFC PV IN gamma synchrony changes depending on whether mice shift to odor or texture-based strategies. **a,** The ratio of gamma synchrony on error vs. correct trials during the pre-decision or post-outcome period for different types of rule shifts. **b,** Gamma synchrony during the pre-decision or post-outcome period of early conflict trials (i.e., conflict trials within the first 5 trials) during different types of rule shifts. **c,** Distribution of mIC-mPFC phase differences (locations of the peak cross-correlation value) during the pre-decision period on error trials (calculated for phase shifts from −360 to 360 degrees). **d,** Comparison of the fraction of in-phase (∼0 degree) vs. anti-phase (∼180 or −180 degree) phase differences during the pre-decision period for texture to odor vs. odor to texture shifts. **e-h,** Comparison of phase differences during post-outcome periods. **e,** Similar to (c) but for the post-outcome period of error trials. **f,** Similar to (d) but for the post-outcome period of error trials. **g,** Similar to (c) but for the post-outcome period of correct trials. **h,** Similar to (d) but for the post-outcome period of correct trials. * p < 0.05. **p<0.01. **** p < 0.0001.

**Supplemental Figure 3:**
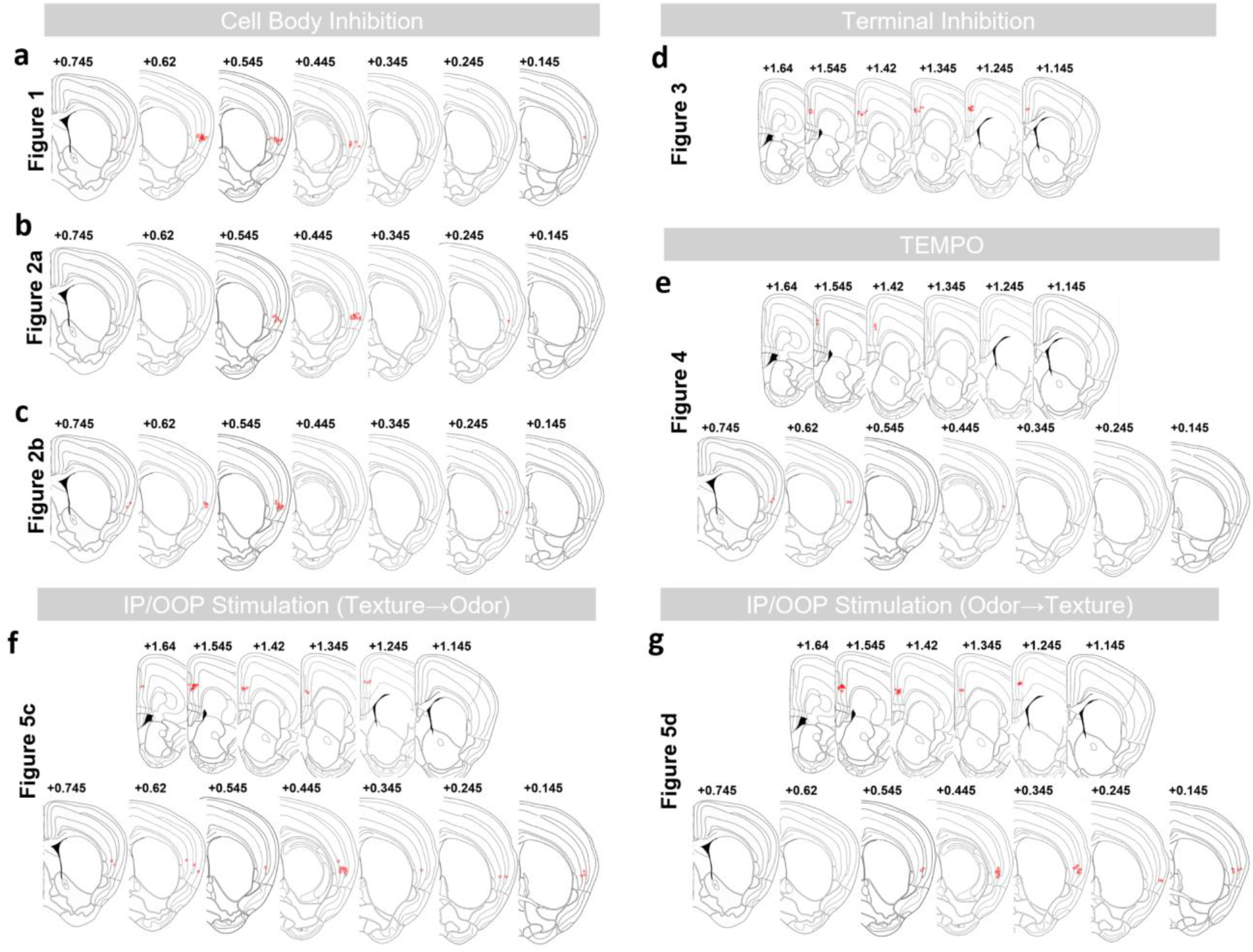
Locations of fiber optic implants. **a-c,** Implant locations within the mIC for cell body inhibition experiments shown in Figures 1 and 2. **d,** Implant locations within the mPFC for mIC to mPFC terminal inhibition experiment shown in Figure 3. **e,** Implant locations within the mIC and mPFC for TEMPO recordings shown in Figure 4. **f-g,** Implant locations within the mIC and mPFC for optogenetic stimulation experiments shown in Figure 5.

**Supplemental Table 1:** Summary of Statistics for Supplemental Figure 2.

| Panel | Time Period | Statistical test |  |  | Multiple comparison |  |
| --- | --- | --- | --- | --- | --- | --- |
| Supplemental Figure 2a | Pre-Decision<br>and Post Outcome | 2-Way ANOVA |  |  | Comparison | P-value |
|  |  | Source of Variation | P value | F (DFn, DFd) | Pre- Decision |  |
|  |  | Interaction | 0.0175 | F (1, 16) = 7.015 | Texture to Odor Shift vs. Odor to Texture Shift | 0.0113 |
|  |  | Phase Shift | 0.255 | F (1, 16) = 1.394 |  |  |
|  |  | Behavior Shift | 0.1816 | F (1, 16) = 1.950 | Post Outcome |  |
|  |  |  |  |  | Texture to Odor Shift vs. Odor to Texture Shift | 0.3891 |
|  |  |  |  |  | Texture to Odor Shift |  |
|  |  |  |  |  | Pre- Decision vs. Post Outcome | 0.0155 |
|  |  |  |  |  | Odor to Texture Shift |  |
|  |  |  |  |  | Pre- Decision vs. Post Outcome | 0.3147 |
| Supplemental Figure 2b | Pre-Decision<br>and Post Outcome | 2-Way ANOVA |  |  | Comparison | P-value |
|  |  | Source of Variation | P value | F (DFn, DFd) | Pre- Decision |  |
|  |  | Interaction | 0.1897 | F (1, 16) = 1.876 | Texture to Odor Shift vs. Odor to Texture Shift | 0.0085 |
|  |  | Phase Shift | 0.0032 | F (1, 16) = 11.99 |  |  |
|  |  | Behavior Shift | 0.0111 | F (1, 16) = 8.246 | Post Outcome |  |
|  |  |  |  |  | Texture to Odor Shift vs. Odor to Texture Shift | 0.304 |
|  |  |  |  |  | Texture to Odor Shift |  |
|  |  |  |  |  | Pre- Decision vs. Post Outcome | 0.0035 |
|  |  |  |  |  | Odor to Texture Shift |  |
|  |  |  |  |  | Pre- Decision vs. Post Outcome | 0.1584 |
| Figure 2d | Pre-Decision Correct Trials | Chi-square |  |  | NA | NA |
|  |  | Comparison | Sides | P value |  |  |
|  |  | In-Phase [-45, 45] vs<br>Anti-Phase [-135,-225] vs<br>Anti-Phase [135,225] | Two | <0.0001 |  |  |
| Figure 2e | Pre-Decision Correct Trials | Chi-square |  |  | NA | NA |
|  |  | Comparison | Sides | P value |  |  |
|  |  | In-Phase [-45, 45] vs<br>Anti-Phase [-135,-225] vs<br>Anti-Phase [135,225] | Two | <0.0001 |  |  |
| Figure 2f | Pre-Decision Correct Trials | Chi-square |  |  | NA | NA |
|  |  | Comparison | Sides | P value |  |  |
|  |  | In-Phase [-45, 45] vs<br>Anti-Phase [-135,-225] vs<br>Anti-Phase [135,225] | Two | <0.0001 |  |  |

## Notes

### Competing Interest Statement

The authors have declared no competing interest.

